# Nighttime gibberellin biosynthesis is influenced by fluctuating environmental conditions and contributes to growth adjustments of Arabidopsis leaves

**DOI:** 10.1101/2020.05.06.080358

**Authors:** Putri Prasetyaningrum, Lorenzo Mariotti, Maria Cristina Valeri, Giacomo Novi, Stijn Dhondt, Dirk Inzé, Pierdomenico Perata, Hans van Veen

**Affiliations:** PlantLab, Institute of Life Sciences, Scuola Superiore Sant’Anna, Pisa 56127, Italy; Dept. of Agriculture, Food and Environment, University of Pisa, Pisa 56124, Italy; Center for Plant Systems Biology, VIB, 9052 Ghent, Belgium; Department of Plant Biotechnology and Bioinformatics, Ghent University, 9052 Ghent, Belgium; Plantecophysiology, Institute of Environmental Biology, Utrecht University, 3584 CH Utrecht

## Abstract

Optimal plant growth performance requires that the action of growth signals, such as gibberellins (GA), are coordinated with the availability of photo-assimilates. Here, we studied the links between gibberellin biosynthesis and carbon availability, and the subsequent effects on growth. The results presented here show that carbon availability, light and dark cues, and the clock ensure the timing and magnitude of gibberellin biosynthesis and that disruption of these mechanisms results in reduced gibberellin levels and expression of downstream genes. Carbon dependent nighttime induction of *GIBBERELLIN 3-BETA-DIOXYGENASE 1* (*GA3ox1)* was severely hampered when preceded by a day of lowered light availability, leading specifically to reduced bioactive GA_4_ levels, and coinciding with a decline in leaf expansion rate during the night. We attribute this decline in leaf expansion mostly to reduced photo-assimilates. However, plants where gibberellin limitation was alleviated had significantly improved expansion demonstrating the relevance of gibberellins in growth control under varying carbon availability. Carbon dependent expression of upstream gibberellin biosynthesis genes (*KAURENE SYNTHASE, KS* and *GIBBERELLIN 20 OXIDASE 1, GA20ox1*) was not translated into metabolite changes within this short timeframe. We propose a model where the extent of nighttime biosynthesis of bioactive GA_4_ by GA3ox1 is determined by starch, as the nighttime carbon source, and so provides day-to-day adjustment of gibberellin responses.

## Introduction

Growth in plants is controlled by many signals, consisting of both environmental and internal cues. Growth is a complex parameter often interpreted as the increase in the number and size of organs (1). Others consider growth the gain of biomass, sometimes more specifically as incorporation of carbon into structural carbohydrates (2-3). Growth parameters like protein and cell wall synthesis directly follow carbon availability (4-5). Carbon storage and utilization are organized such that carbohydrates are roughly evenly incorporated into structural biomass throughout the day (2-3). However, size increases of plants do not necessarily follow carbon availability. Hypocotyl and leaf expansion rates have been shown to vary throughout the day, and these expansion patterns have been attributed to the circadian clock and light signalling (6-8). Interestingly, expansion and metabolic biosynthesis can be distinctly regulated. In hypocotyls, cell wall biosynthesis was exclusively depended on metabolic signals and cell expansion on the circadian clock (5). For optimal plant performance, size increase must match biomass integration. The ability to adjust leaf expansion rates based on resource availability would be especially important when conditions vary from day to day.

In conjunction with the clock and light signals, a variety of growth stimulating signalling networks exists within the plant, including a set that rely on phytohormones (1). Pivotal among these are gibberellins (GAs), which positively regulate cell expansion and cell division (9-10). Plants deficient in gibberellins develop slowly and are typically very small in stature (11), whereas overexpression of biosynthetic routes or gibberellin signalling leads to enhanced plant size (12-15). Gibberellin biosynthesis occurs via a sequence of enzymatic steps, starting in the plastids and subsequently the endoplasmic reticulum, which convert geranylgeranyl diphosphate to GA_12_. GA_12_ is considered the common precursor to all gibberellins in plants and is further processed into a variety of forms by GIBBERELLIN 20-OXIDASEs (GA20ox). The final step to bioactive gibberellin is catalysed by GIBBERELLIN 3-BETA-DIOXYGENASE (GA3ox) (11). In higher plants GA_4_, GA_1_ and GA_3_ have been confirmed to bind the GIBBERELLIN INSENSTIVE DWARF1 (GID1) receptor (16-17). These bioactive gibberellins can also be made inactive by the enzyme GA2ox (11).

Established aspects of the regulation of gibberellin biosynthesis concern localized suppression in the shoot apical meristem to ensure stem cell maintenance, and feedback regulation that is considered important to maintain gibberellin homeostasis (18-19). Feedback regulation is structured such that it aims to keep gibberellin levels constant (19) and is therefore not adequate to explain growth adjustments to varying conditions. To identify the role of gibberellins in vegetative growth and adjustment to adverse conditions we studied the regulation of key gibberellin biosynthetic genes, the subsequent variation in gibberellin abundance and ultimately their role in growth in *Arabidopsis thaliana*. This led to the finding that nighttime conversion of precursors to bioactive GA_4_ by the starvation sensitive *GA3ox1*, plays a role in adjusting growth to adverse conditions.

## Results

### Light, the circadian clock and carbon availability regulate the timing and transcriptional induction of GA biosynthetic enzymes

Several enzymatic steps of GA biosynthesis are encoded by multigene families. To identify the key family members with relatively high expression during vegetative growth, two public microarray datasets that followed transcript abundance throughout 48 hours in adult Arabidopsis rosettes were investigated (20-21). Further indication of a role in adjusting growth control was based on oscillatory behaviour over the day-night cycle, indicating a responsiveness to changes in carbon supply, light signals or the clock. These data (Suppl. Figure 1) pointed to *KAURENE SYNTHASE (KS), GA20ox1* and *GA3ox1* as main candidates (Figure 1).

**Figure 1.**
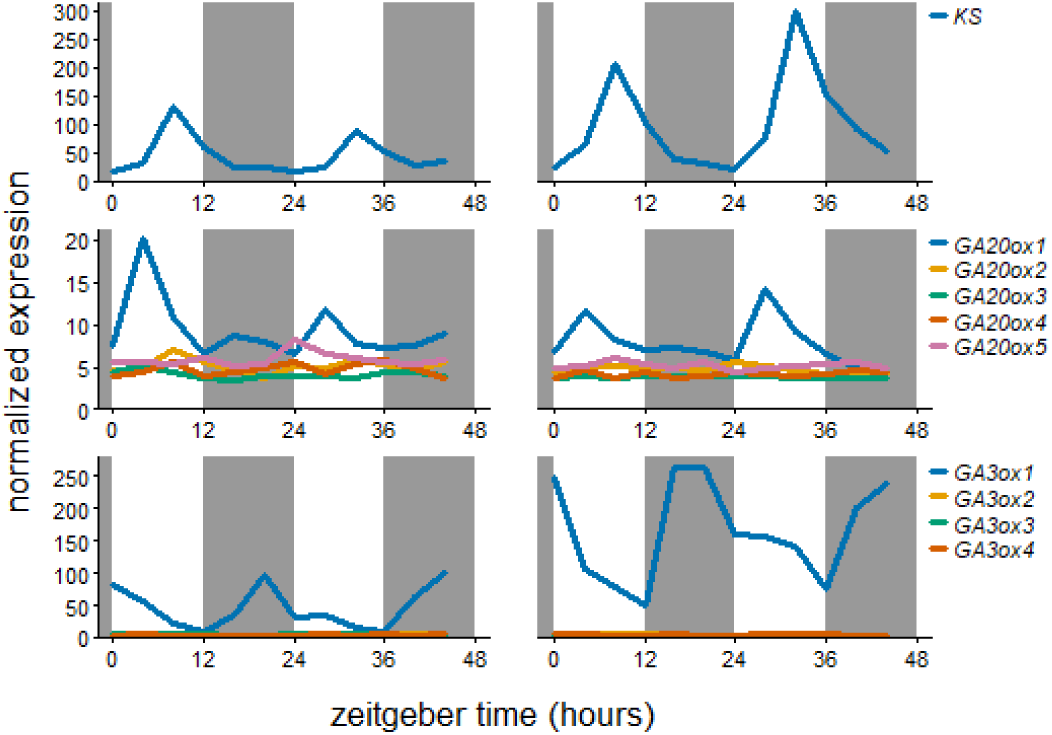
mRNA abundance over the day-night cycle of the gene family members of rhythmic GA biosynthetic genes. Left and right column based on data from 35 day and 29 day old, respectively, soil grown Col-0 rosettes (20-21). Grey boxes represent the night period. All family members are shown.

Clock components and clock-regulated genes retain their oscillations after a transfer to continuous light conditions (Supplementary Figure S2A). Of the daytime expressed candidates, oscillations were retained for *KS*, but not for *GA20ox1*. However, their peak in expression during normal day/night cycles, start of the day (*GA20ox1*) and afternoon (*KS*), matched the publicly available dataset (Figure 2A). *GA20ox1* oscillations seemed independent of the clock (Figure 2A), and a role for light in inducing this gene was investigated by exposing plants to light three hours prior to expected dawn. An earlier start of the day resulted in a concomitant earlier peak of *GA20ox1* (Figure 2B). Since an early day also led to a sooner increase in sugars (Figure 2B, Supplementary Figure S2B), a role for photosynthates to induce the earlier peak could not be excluded. After inhibiting photosynthesis with DCMU, *GA20ox1* still peaked upon earlier light exposure. However, DCMU did reduce the extent of induction and abolished *GA20ox1* induction at later timepoints (Figure 2B). Carbon starvation reduced *GA20ox1* expression, but a DCMU treatment can be very severe, as apparent from the very strong induction of carbon starvation marker genes, *DIN6* and *TPS8* (Supplemental Figure S2B). Therefore, plants were also exposed to low light conditions that were sufficient to maintain a positive carbon balance but did reduce photosynthesis and also showed minor induction of starvation markers compared to the DCMU treatment (Supplementary Figure S2C-E). Both *KS* and *GA20ox1* showed severely reduced daytime expression after DCMU treatment, whereas the low light treatment led, as expected, to an intermediate reduction in expression levels (Figure 2C).

**Figure 2.**
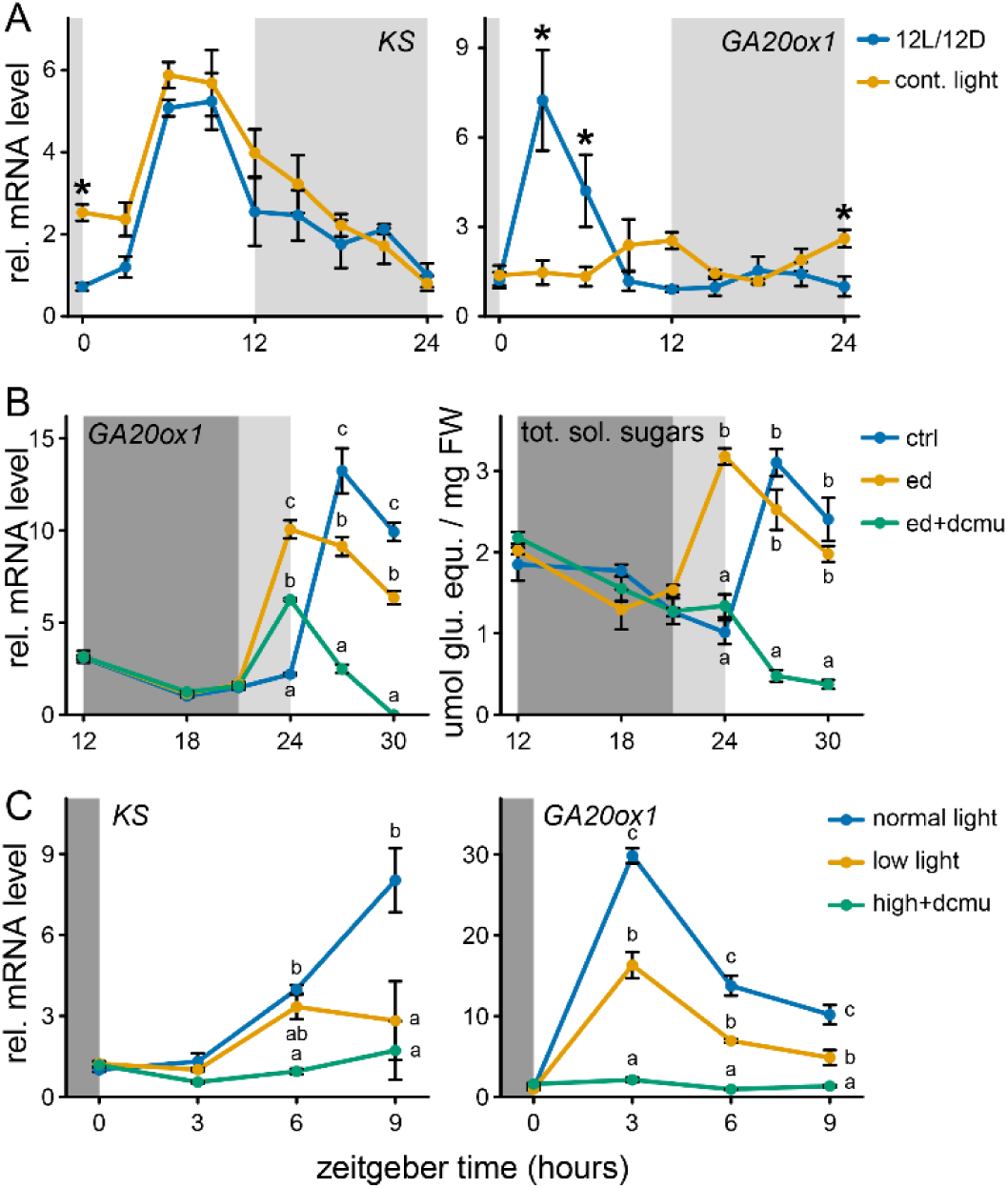
Transcriptional regulation of the daytime peaking GA biosynthetic genes, *KS* and *GA20ox1*. **A**. The role of the circadian clock in mediating rhythmic expression. Light grey boxes represent subjective night. Sampling started the second day after beginning continuous light conditions. **B**. Effect of a 3 hour early start of day. Dark and light grey represent night and the subjective night, respectively. DCMU and mock solution (100 µM) was applied at ZT12. Ctrl: 12L/12D, ed: early start of day. **C**. The effect of reduced light availability or DCMU on KS and *GA20ox1* transcript abundance. DCMU was applied at the start of the preceding night. All data are from ∼10 leaf stage soil grown Col-0 rosette. Mean ± sem are shown, n=4. Asterisks (planned comparisons) and letters (Tukey HSD) represent statistically significant difference at specific time points p < 0.05)

Regarding the nighttime expressed candidate, *GA3ox1* induction was induced upon transition to darkness, in line with public data (Figure 1 and 3A). Oscillations of *GA3ox1* were not retained under continuous light, indicating that a shift to darkness is essential for a *GA3ox1* induction (Figure 3A). However, starting the night period earlier or later did not lead to a concomitant shift in *GA3ox1* induction (Figure 3B and C). This implies that *GA3ox1* responds to darkness, but only when it occurs simultaneously with the expected onset of the night. The effect of carbon starvation on the nighttime *GA3ox1* induction was investigated by DCMU and low light treatments, like done for daytime expressed *KS* and *GA20ox1*. Although low light treatments led to an intermediate starvation response compared to DCMU (Supplemental Figure 2E), it was sufficient to completely abolish the nighttime induction of *GA3ox1*, identical to the effect of DCMU (Figure 3D). These observations suggest that *GA3ox1* is more sensitive to carbon starvation. The relevance of energy and carbon signalling for *GA3ox1* regulation was further investigated by transgenics, mutants and pharmaceutical inhibitors of three main energy signalling pathways, TARGET OF RAPAMYCIN (TOR) kinase, SNF1 RELATED PROTEIN KINASE 1 (SnRK1.1/KIN10) and glucose signalling (GLUCOSE INSENSITIVE2, GIN2)(22-24). Overexpression of the wild type alleles of neither KIN10 or TOR, nor the *gin2-1* null allele affects the behaviour of *GA3ox1* during night. However, preventing the activity of KIN10 and TOR kinase by either an active site specific mutation or pharmaceutical inhibitors, led to an enhanced or abolished nighttime *GA3ox1* induction, respectively (Supplemental Figure 3). The direction of change corresponded to the removal of either a starvation, KIN10, or an energy abundance signal, TOR.

**Figure 3.**
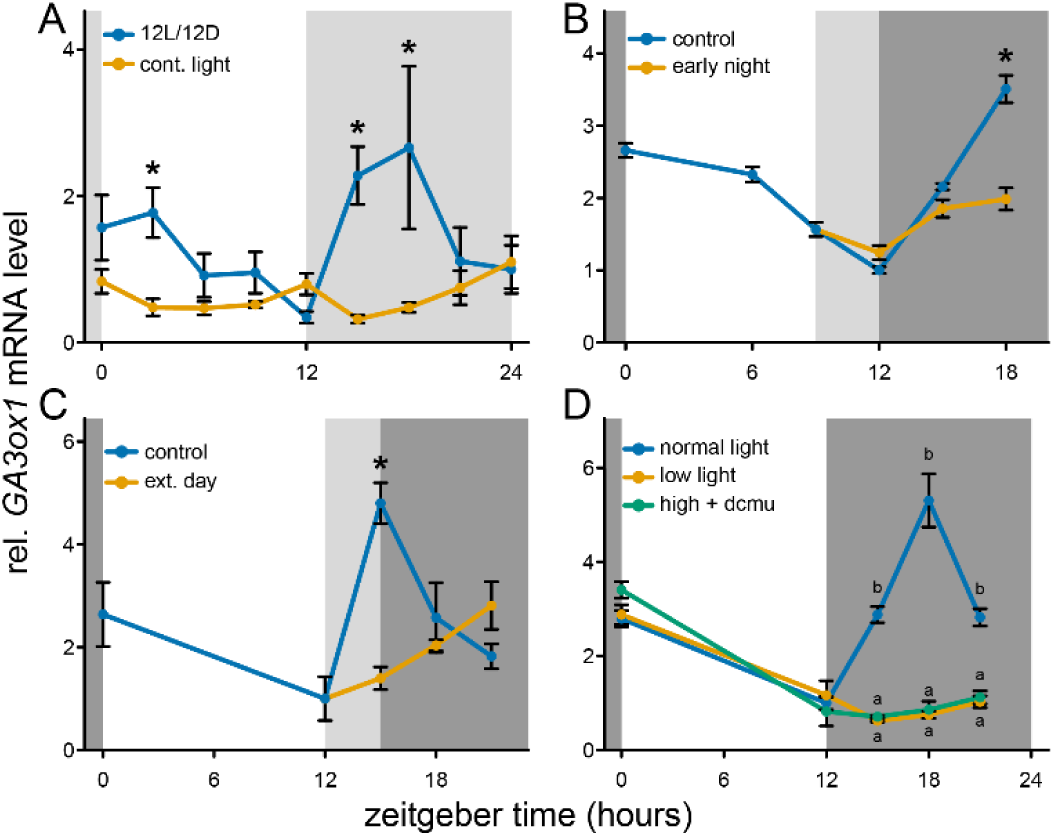
Transcriptional regulation of the night-time peaking GA biosynthetic gene, *GA3ox1*. **A**. The role of the circadian clock in mediating rhythmic expression. Light grey boxes represent subjective night. Sampling started second day after starting continuous light conditions. **B**. Effect of a 3 hour early start of the night. Dark and light grey represent night and the subjective night, respectively. **C**. Effect of a 3 hour extension of the day. Dark and light grey represent night and the subjective night, respectively. **D**. The effect of reduced light availability or DCMU on nighttime *GA3ox1* transcript abundance. DCMU was applied at the start of the preceding night. All data are from ∼10 leaf stage soil grown Col-0 rosette. Mean ± sem are shown, n=4. Asterisks (planned comparisons) and letters (Tukey HSD) represent statistically significant difference at specific time points p < 0.05)

### Bioactive gibberellins accumulate during the night and are severely reduced the morning after a day of low light levels

Patterns of transcriptional regulation suggest that gibberellin precursors are made during the day via the upstream enzyme *KS* and pre-final enzyme *GA20ox1*. These accumulated precursors would then be converted to bioactive GA by *GA3ox1* which peaks during the night, leading to high bioactive GA at the end of the night. Moreover, reduced light availability affected all three transcripts, especially *GA3ox1* (Figure 2C and 3D). To investigate whether such transcriptional patterns would be translated into changes in the corresponding compounds, we measured gibberellin content at the start and end of the day. Moreover, we confronted plants with an extended day to reduce *GA3ox1* induction whilst maintaining carbon availability (Figure 3C).

Though the gibberellin precursor pool (GA9, GA19 and GA20) appeared higher in the evening, no significant differences were observed between timepoints nor between light levels (Figure 4A). The bioactive gibberellins differ in their affinity for the receptor GID1, with GA_3_ and GA_1_ having a weak affinity compared to strong binding affinity of GA_4_ (16-17). The low affinity bioactive gibberellins (GA_1_ and GA_3_) had no clear differences between treatments (Supplemental Figure 4). The high affinity gibberellin, GA_4_, was the most abundant bioactive gibberellin, and its levels were highest at the end of the night (Figure 4B). Reducing nighttime *GA3ox1* induction by an extended day (Figure 3C), led to a subsequent drop in GA_4_ the following morning (Figure 4B). Low light levels led to an even stronger drop in GA_4_ the following morning, whereas GA_4_ levels remained unchanged at the end of the actual low light day (Figure 4B). These results suggest an important role of nighttime *GA3ox1* expression in determining bioactive GA levels in response to low light, whereas the drop of *KS* and *GA20ox1* was not translated into reduced gibberellin biosynthesis within the time frame studied here. Gibberellin inactivating enzymes could also play a role in determining the dynamics of bioactive GA. However, no consistent changes of inactivated GAs (GA_8_, GA_29_ and GA_34_) were found (Supplemental Figure 4).

**Figure 4.**
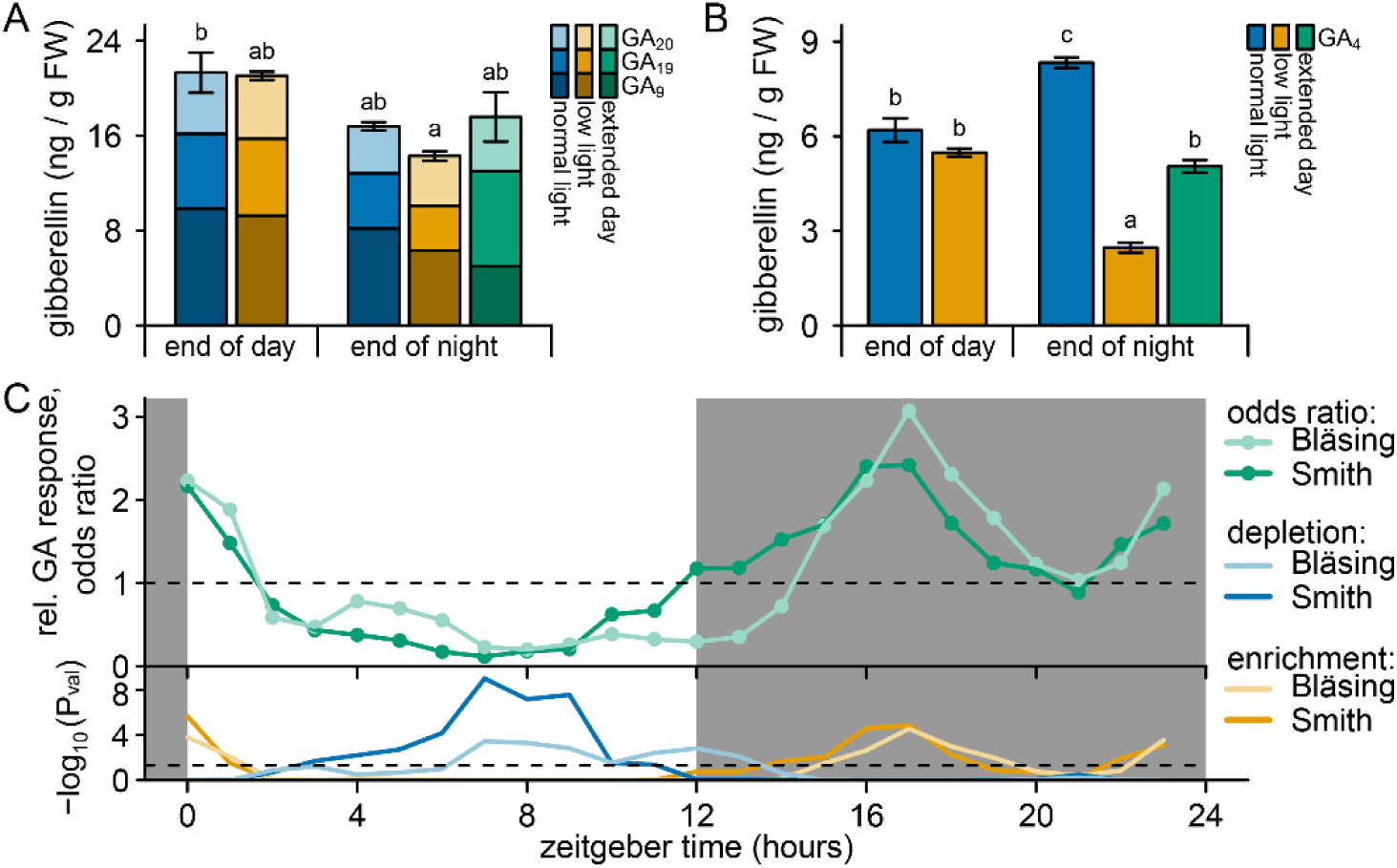
GA abundance and responses over time and after low light conditions. **A**. Abundance of gibberellin precursors metabolized by GA20ox, directly following a control (high light) or low light day (end of day, ZT12) and after the following subsequent night (end of night, ZT24). Extended day (under control light conditions) represents a 3 hour delay in the start of the night as in Figure 2C. Data are mean, sem of the sum of precursors is shown. Letters indicate significant differences (P < 0.05, Tukey HSD, N=3). **B**. Conditions as in A, but levels of the main bioactive GA4 are shown. **C**. Relative enrichment (odd score > 1) or depletion (odd score < 1) of GA responsive genes among gene-sets that have rhythmic behaviour peaking at specific times of the day. Gene sets for each time of day were obtained from (20-21, 47). GA responsive genes were based on GA treated rosettes (22). Below is the statistical significance of depletion or enrichment of GA responsive genes (Hypergeometric distribution). Dashed line indicates P = 0.05.

Lower and higher levels of GA_4_ at the end of the day and night, respectively, suggest temporal dynamics in GA abundance. These direct gibberellin measurements are supported by *in silico* investigation of genes responsive to GA in Arabidopsis rosettes (25). For two independent datasets (20-21), the GA responsive genes were enriched among gene-sets that peaked in expression during the night, but underrepresented (depleted) among daytime peaking genes (Figure 4C, Suppl. Figure 5). These dynamics of gibberellin responsive genes could also be ascribed to changes in gibberellin sensitivity, as also observed in seedling hypocotyls (26). Indeed, transcript levels of *GID1A* seem to be highest during the evening and start of the night in rosettes (Supplemental Figure 1). The starchless mutant *phosphoglucomutase* (*pgm*), typically suffers from severe starvation at night (21,27). Subsequently, the nighttime induction of *GA3ox1* was abolished, whereas the daytime GA biosynthetic genes (*KS* and *GA20ox1*) remained mostly unaffected (Supplemental Figure 6A). Indeed, *pgm* and other starch mutants are susceptible to altered GA metabolism and reduced gibberellin levels (28). Subsequently, we found that rhythmic expression of GA responsive genes was lost in *pgm*, despite retaining rhythmicity in *GID1A* (Supplemental Figure 6B). The relevance of gibberellins in time of day specific leaf expansion was further investigated in the *gibberellic acid insensitive* (*gai*) and *della pentuple* (*pent*) mutants, which cannot respond to varying gibberellin levels (15). Expansion rates of *gai* and *pent* were decreased at night, when the expression of *GA3ox1* determines the bioactive GA levels (Figure 5). Additionally. Expansion rates were elevated in the morning (Figure 5), which might represent a GA-independent behaviour that might utilize untapped nighttime growth potential. Likewise, a GA3ox1 knockout mutant, *ga3ox1-3* (29), also has reduced nighttime expansion and elevated morning expansion (Figure 5). Overall, this suggests that an ability to respond to changes in gibberellin levels and GA biosynthesis contributes to time of day specific expansion.

**Figure 5.**
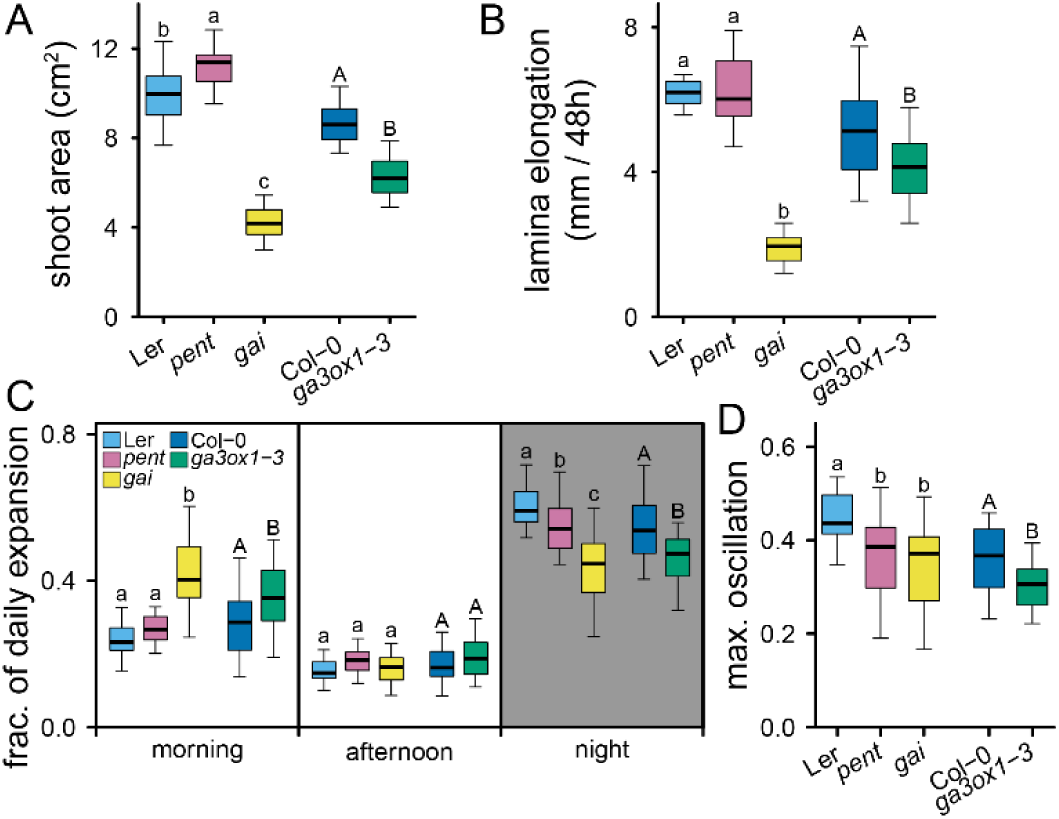
Daily changes in expansion. **A**. Projected shoot area 3 weeks after transplanting. **B**. Lamina expansion of leaf 7 in a 10 leaf rosette over 48 hours. **C**. portion of the daily expansion occurring during the morning (ZT0 – ZT6), afternoon (ZT6-ZT12) and night (ZT12-ZT24). The average of two days is shown. **D**. The extent of growth rhythmicity expressed as the maximum difference in fraction of daily expansion between the three time periods investigated (morning, afternoon and night) growth rate difference. Letters indicate significant differences (P < 0.05, Tukey HSD, N=25), lower and uppercase distinguish comparisons for the two different genetic backgrounds used. *Pent* = *della pentuple* mutant.

**Figure 6.**
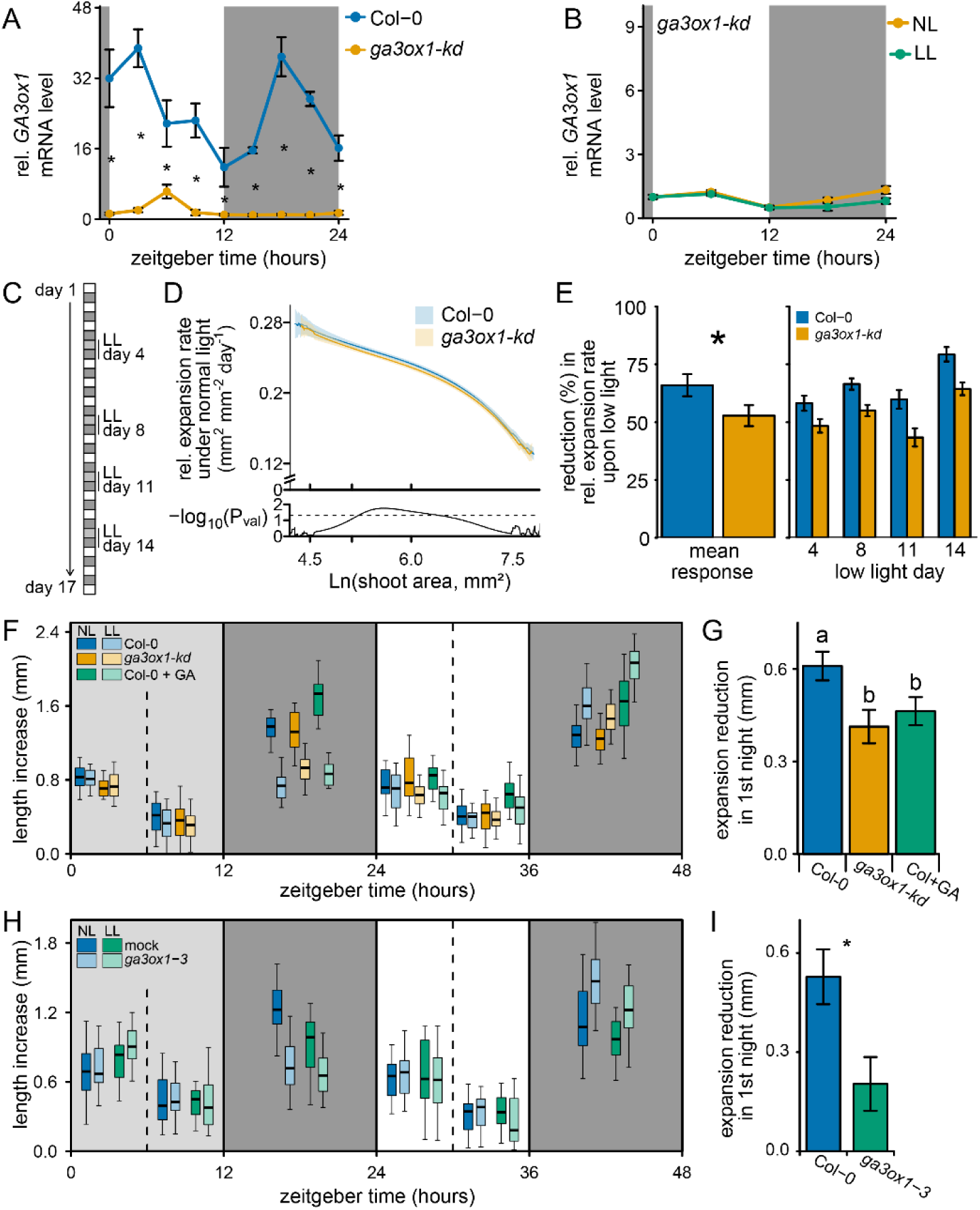
Carbon and GA dependent growth under constant and variable conditions. **A**. Transcript levels of *GA3ox1* in Col-0 and the *ga3ox1-kd* line, mean ± sem (N=4). **B**. The behaviour of *GA3ox1* during low light (LL) and normal light (NL) conditions in *ga3ox1-kd*, mean ± sem (N=5). **C**. A brief schematic of the measuring and treatment regime. **D**. Expansion rate during constant high light conditions for a given projected shoot area. The shaded areas indicate the 95 % confidence interval of the mean. The statistical difference between genotypes (students t-test) is given below (N=40). Dashed line indicates P = 0.05. Based on control plants (NL) not confronted by low light days. **E**. The reduction in relative expansion rate during a low light day and its subsequent night compared to the expansion rate of the previous days. The reduction for each individual low light day, and the mean low light response over all 4 days are shown. Asterisks indicate the significant difference (ANOVA), mean ± sem (N=40). **F**. Lamina length increase (leaf 7) over measured time intervals (6 or 12 hours), during and following a low light day. Dark and light grey boxes indicate the night and the low light treatment respectively. Gibberellin (GA4+7)/mock was applied at ZT12. N=25, 10 leaf stage plants. **G**. Reduction in expansion compared to normal light plants specifically at the night following a low light treatment (ZT12 to ZT24), mean ± sem (N=25). Low light Col+GA growth reduction is compared to Col-0 normal light + mock. Letters indicate significant differences (P < 0.05, Tukey HSD). **H** and **I**. Experiment identical to F and G, but with Col-0 and the full knock out *ga3ox1-3*. Mean ± sem (N=25) is shown, asterisk indicates statistical significance. NL: control normal light conditions, LL: low light conditions.

### Reduced light availability leads to reduced leaf expansion rates during the following night only, which is partially mediated by *GA3ox1*

The next step was to investigate whether the carbon dependent regulation of *GA3ox1* transcription and subsequent GA_4_ levels are important to adjust growth to periodic low light levels. To effectively investigate growth reductions, it is important that a mutant only has minor growth effects under control conditions (termed normal light), therefore a *ga3ox1* knock down (SALK_025076) with a T-DNA insertion in the intron was used. This *ga3ox1-kd* had a more than 10 fold drop in *GA3ox1-kd* mRNA abundance and less than half the GA_4_ levels (Figure 6A, Supplemental Figure 7A-C). Furthermore, the low *GA3ox1-kd* mRNA abundance of the knock down was not significantly further reduced by a low light treatment (Figure 5B). The specific leaf area, an important growth parameter, was unaffected. However, *ga3ox1-kd* did flower slightly later and had reduced bolt length (Supplemental Figure 7D).

With the automated phenotyping platform WIWAM XY to precisely control soil water content and to image individual plants over time (30,31) we followed the expansion rates of Col-0 and *ga3ox1-kd* plants under constant day/night cycles and when confronted with four interspersed low light days (Figure 6C). To obtain expansion rates under normal light and to determine low light mediated reductions in expansion we fitted growth curves to individual plants (Supplemental Figure 8). Under control conditions expansion rates were close to identical between Col-0 and *ga3ox1-kd* (Figure 6D). We concluded that GA3ox1-dependent fluctuations in gibberellin levels could not be the main driver of growth in *ga3ox1* under normal light conditions. Exposure to the four low light days led to strongly reduced final rosette area in both genotypes (Supplemental Figure 8), implying that a few low light days can have dramatic effects. Quantification of the reduction in expansion during the low light day and subsequent night showed that *ga3ox1-kd* had a smaller reduction in expansion rate than Col-0 (Figure 6E). This suggests that expansion in *ga3ox1-kd*, in which the nighttime expression and regulation of *GA3ox1* is negligible (Figure 6A-B), does not suffer as strongly from low light as the wild type, where low light severely dampens the expression of *GA3ox1* (Figure 3D) and GA_4_ levels (Figure 4B).

To investigate the exact time of the day at which the growth reduction takes place and to further explore the role of GA3ox1 in such reductions, leaf expansion rates of the fastest growing leaf (7^th^ leaf; Supplemental Figure was followed over 6- or 12-hour intervals. During the low light treatment, leaf expansion rates remained unchanged, whereas in the subsequent night, expansion was severely reduced (Figure 6F). Preventing a drop in GA_4_ after low light by either gibberellin application or the *ga3ox1*-kd, mitigated the reduction in expansion to a similar degree (Figure 6G). However, neither manipulation of gibberellins could completely abolish the growth penalty (Figure 6G). Similarly, for the full knock out of *GA3ox1, ga3ox1-3*, low light induced reduction of nighttime leaf expansion was mitigated (Figure 7H-I). Low light levels also lead to altered carbon dynamics in the subsequent day. The following day less carbon is allocated to structural biomass and subsequently soluble carbohydrates (including starch) accumulate to higher levels (32), and this accumulation the subsequent day was independent of gibberellins (Supplemental Figure 10A-B). The higher carbon availability coincided with enhanced growth during the second night after the low light treatment, especially in Col-0 where the low–light-induced expansion was of a similar magnitude as the expansion induced solely by exogenous gibberellins. The *ga3ox1-kd* seemed not able to benefit from this growth increase to the same extent (Supplemental Figure 10C).

## Discussion

A capacity to adjust hormonal profiles to the prevailing environmental conditions is essential for plants to ensure optimal performance. For this reason, we aimed to identify how gibberellin metabolism was affected by carbon availability and the subsequent effects on growth. Nighttime production of GA_4_ by GA3ox1 was found to be highly sensitive to low light levels, which led to a reduction of GA_4_ levels only in the subsequent night and not the actual day. Similarly, effects on leaf expansion upon low light were only noticeable during the night and were mitigated in plants whose growth was not driven by nighttime expression of GA3ox1. The results presented here provide new insights into the regulation of gibberellin metabolism and add perspective on the mechanisms of growth control.

We found that reducing *GA3ox1* mRNA levels, either through an extended day, low light availability or a genetic knockdown, consistently led to reduced GA_4_ abundance, but did not affect the levels of the low affinity bioactive gibberellins (GA_3_ and GA_1_, Figure 4). Indeed, *in vitro* activity of GA3ox1 was shown to have a strong preference for converting precursors to GA_4_ (33). Full knockouts of GA3ox1 were also reported to have reduced GA_4_ levels (29,34). In contrast, clearly reduced mRNA levels of upstream biosynthesis genes (*KS* and *GA20ox1*, Figure 2) did not lead to a corresponding reduction of precursors (Figure 4). GA20ox is generally considered the rate limiting step, since constitutive overexpression leads to higher GA_4_ levels, in contrast to overexpression of other biosynthesis genes (18). Longevity of prior produced enzymes could explain a lack of responses at the metabolite level observed after the short transcriptional perturbation of this study. Likely, successive days of reduced light availability would eventually also lead to an effect on precursor metabolism. However, GA_4_ levels were quick to respond to changes in *GA3ox1*, suggesting that GA3ox1 proteins need to be resynthesized daily. This implies that GA metabolism is reset every day via *GA3ox1*. The results presented here suggest that the abundance of GA3ox1 and GA20ox1 vary throughout the day-night cycle, and subsequently also the flux through each of these pathways.

It is tempting to attribute the low afternoon-growth and high night- and morning-growth (Figure 5C and 6F) with the corresponding changes in GA_4_ levels and behaviour of GA_4_ responsive genes (Figure 4). Gibberellin sensitivity was shown to be regulated by the circadian clock (26), here we show that also the actual levels of GA_4_ vary throughout the day and probably coincide with high gibberellin sensitivity (Figure 4, Supplemental Figure 1). Varying expansion rates of leaves throughout the day-night cycle are also regulated by carbon availability, water availability, light cues and the circadian clock (6-8, 31, 35-36). It is not straight forward to piece apart the contributions of these growth cues and discern how they act in concert. Nevertheless, the results presented here show that light and dark cues, and the clock ensure timing of gibberellin biosynthesis and that disruption of biosynthesis patterns in the starchless *pgm* abolishes the rhythmic nature of downstream GA responses. Moreover, removing the ability to respond to varying gibberellin levels or disrupting biosynthesis impacts growth dynamics corresponding to the rhythmicity in GA biosynthesis.

The nighttime induction of *GA3ox1* is highly dependent on carbon availability (Figure 3D). The consistently reduced nighttime growth penalty upon low light, either for whole rosettes or a single leaf, of plants without the capacity of *GA3ox1* regulation demonstrates the relevance of this gene and subsequent gibberellins in adapting growth to the prevailing conditions (Figure 6). Although, the difference in the low light induced growth penalty between Col-0 and *ga3ox1-kd* is seemingly small, around 20-30 %, it is of a similar magnitude as a rescue by gibberellins (Figure 6E and G). A variety of growth stimulating gene regulatory networks have been identified in Arabidopsis and many of these act independent of one another (12-13). Likewise, the different plant hormones have surprisingly little overlap in downstream genes and appear to operate in distinct fashions (37). Alternative mechanisms that connect carbon and growth networks likely exist in plants (38). One is the starvation induced autophagy of brassinosteroid signalling components (39,40). A collection of numerous of such growth regulating mechanisms will buffer against the loss of the single mechanism identified here, which could explain the small effect size. Similarly, boosting carbon assimilation by elevated CO_2_ can still enhance growth in paclobutrazol treated plants (41). In this study, plants were confronted by fewer photo-assimilates, which will place physical constraints on growth regardless of the manipulation of growth signalling mechanisms.

Gibberellins are crucial hormones for plants, which stimulate growth and development. This study reveals how gibberellin metabolism is balanced to match the physiology of the plant. We present a model of gibberellin biosynthesis that is coordinated throughout the day-night cycle by light cues, dark cues and the circadian clock. Here, the nighttime GA_4_ biosynthesis rate is reset daily and strongly depends on carbon availability. Starch, as the nighttime carbon source (42), plays a pivotal role in this model and acts as a robust integrator of daytime performance, unlike the highly variable nature of photosynthesis and light availability. Using starch, rather than photosynthesis directly, could prevent excessive starting and stopping of growth. Though the exact mechanism is unclear, plants can precisely estimate starch levels to ensure proper nighttime utilization rates (42,43). Subsequently, starch utilization rates could be accurately used to pace gibberellin metabolism and aid in adjusting gibberellin responses, such as growth, to the prevailing conditions when plants grow in variable environments.

## Methods

### Plant material, growth and treatments

Col-0 and *ga3ox1* (NASC: N670439) were grown on a soil-perlite mixture. Light levels used for the experiments ranged from 100-130 µmol m^-2^ s^-1^ photosynthetic active radiation (PAR) and relative humidity between 50 and 70 % and a temperature of 20°C. Low light treatments were between 30-45 µmol m^-2^ s^-1^ PAR. DCMU (100 µM) and GA_4+7_ (100 µM) were applied by spraying on the plants. Experiments were started once the plants had 11 visible leaves.

### Gene expression and carbohydrates

For Gene expression two entire rosettes were pooled per replicate. RNA isolation was performed according to Kiefer method (40). DNA contaminant was removed with RQ1 RNase-Free DNase™ (Promega) and cDNA was synthesized using Maxima First Strand cDNA Synthesis Kit™ (Thermofisher). The cDNA was used as template for real-time PCR (primers in Supplemental table 1) using CFV384 Touch™ Real-Time PCR Detection System (Bio-Rad). Housekeeping genes (*AT4G34270, AT1G13320*) were selected based on ref. 45, with an extra focus on constants expression levels during the day-night cycle. Sugars and starch were determined through coupled enzymatic reactions and NADH absorbance at 340 nm as described previously (28).

### Gibberellin metabolites

Gibberellin levels were determined as previously described in ref. 46 with some modifications. In short, 2 to 4 grams of pooled shoot was homogenized in cold 80% (v/v) methanol (1:5, w/v) using a mortar and pestle. Fifty nanograms of deuterated GAs ([17,17-2H2]-GA9, [17,17-2H2]-GA4, [17,17-2H2]-GA34, [17,17-2H2]-GA19, [17,17-2H2]-GA20, [17,17-2H2]-GA29, [17,17-2H2]-GA1, [17,17-2H2]-GA8, [17,17-2H2]-GA3) were added as internal standards to account for purification losses. Methanol was evaporated under a vacuum at 35°C, and the aqueous phase was partitioned against ethyl acetate, after adjusting the pH to 2.8. The extracts were dried and suspended in 0.3-0.5 ml of distilled water with 0.01% acetic acid and 10% methanol. HPLC analysis was performed with a Kontron instrument (Munich, Germany) equipped with a UV absorbance detector operating at 214 nm. The samples were applied before to a 150 × 4.6 mm ID column packed with ODS Hypersil (Thermo Fischer Scientific, Milan Italy), particle size 5 μm, eluted at a flow rate of 1 ml min^-1^. The column was held constant at 10% methanol for 4 min, followed by a double gradient elution from 10% to 30% and 30% to 100% over 40 min. The fractions corresponding to the elution volumes of standard GAs were collected separately. Subsequently, the fractions were applied to 250 × 4.6 mm ID, Nucleosil 100-5 N(CH3)2 column (Macherey-Nagel GmbH & Co, Düren, Germany) and eluted isocratically with 100% methanol containing 0.01% acetic acid at a flow rate of 1 ml min^-1^. The fractions corresponding to the elution volumes of standard GAs were collected, dried and silylated with N,O bis (trimethylsilyl) trifluoroacetamide containing 1% trimethylchlorosilane (Pierce, Rockford, IL, USA) at 70 °C for 1-h. The gas chromatography–tandem mass spectrometry analysis was performed on a Saturn 2200 quadruple ion trap mass spectrometer coupled to a CP-3800 gas chromatograph (Varian Analytical Instruments) equipped with a MEGA (http://www.mega.mi.it) 1MS capillary column (30 m 3 0.25-mm i.d. and 0.25-µm film thickness). The carrier gas was helium, which was dried and air free, with a linear speed of 60 cm s^-1^. The oven temperature was maintained at 80 °C for 2 min and increased to 300 °C at a rate of 10 °C min^-1^. Injector and transfer line were set at 250 °C and the ion source temperature at 200 °C. Full scan mass spectra were obtained in EI+ mode with an emission current of 10 µA and an axial modulation of 4 V. Data acquisition was from 150 to 650 Da at a speed of 1.4 scan s^-1^. GAs were identified by comparing the full mass spectra with those of the authentic compounds, and quantification with reference to standard plots of concentration ratios versus ion ratios that were obtained by analysing known mixtures of unlabelled and labelled GAs.

### Growth measurements

The Weighing, Imaging and Watering Automated Machine (WIWAM) (www.wiwam.com) to precisely control soil water content and image plants was used to follow size increase of individuals over time. The PSB Interface for Plant Phenotype Analysis (PIPPA, https://pippa.psb.ugent.be) was used for analysis, visualization and management of phenotypic datasets and images. A four-parameter logistic model (constant conditions) or a series of exponential models (low light treatment) were fitted to individual plants. This allowed us to track growth rates over time and rosette area. Contributions of distinct leaves to total size increase was determined by two destructive harvests, separated by 24 hours. Leaves were dissected and arranged on 0.59% agarose filled plates, leaf incisions ensured proper flattening of the leaves. Areas were determined with ImageJ software on pictures obtained by scanning the plates. Growth of individual leaves over time were determined, non-destructively, by manual length measurements with a digital calliper.

### In silico analysis and statistics

Data analysis and statistics were done using R software. Data were transformed if needed to satisfy the assumptions of statistical tests. Calling of rhythmic genes in *pgm* was done by the HAYSTACK tool from the DIURNAL project (47).

## Acknowledgments

This work was supported by the Scuola Superiore Sant’ Anna (SSSA) and the Plantecophysiology group (Utrecht University). P.Pr was supported by a PhD fellowship in Agrobiodiversity (SSSA). P.Pe. and L.M. acknowledge support of CISUP at University of Pisa. S.D. was funded by Research Foundation Flanders (FWO) by a Postdoctoral Fellowship. H.v.V was supported by SSSA and grant ALWOP.419 from the dutch scientific organisation. We further acknowledge financial support by the Access to Research Infrastructures activity in the Horizon2020 Programme of the EU (EPPN2020 Grant Agreement 731013).

## Notes

### Competing Interest Statement

The authors have declared no competing interest.

