## SupplementalMaterial for "Nighttime gibberellin biosynthesis is influenced by fluctuating environmental conditions and contributes to growth adjustments of Arabidopsis leaves"

Author contributions: P.Pe and H.v.V conceived the project; P.Pr, S.D., D.I. P.Pe and H.v.V designed experiments; P.Pr, L.M., M.C.V., G.N. and H.v.V Performed Experiments; P.Pr, S.D. and H.v.V analysed the data; H.v.V. and P.Pe wrote the paper with contributions from S.D. D.I.

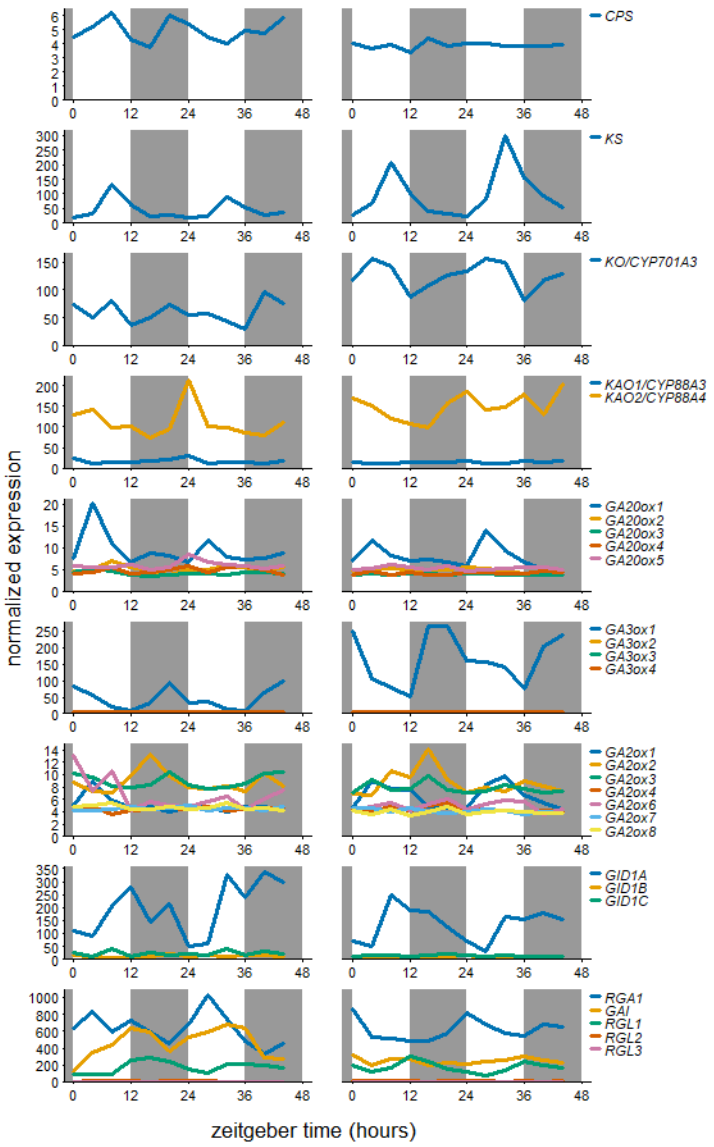


**Supplemental Figure 1. mRNA abundance over the day-night cycle of GA biosynthetic and signalling genes.**

Left and right column based on public data from 35 day and 29 day old, respectively, soil grown Col-0 rosettes at light intensity of 130 µmol $m^{-2}$ $s^{-1}$ (20-21). Grey boxes represent the night period. All family members are shown.

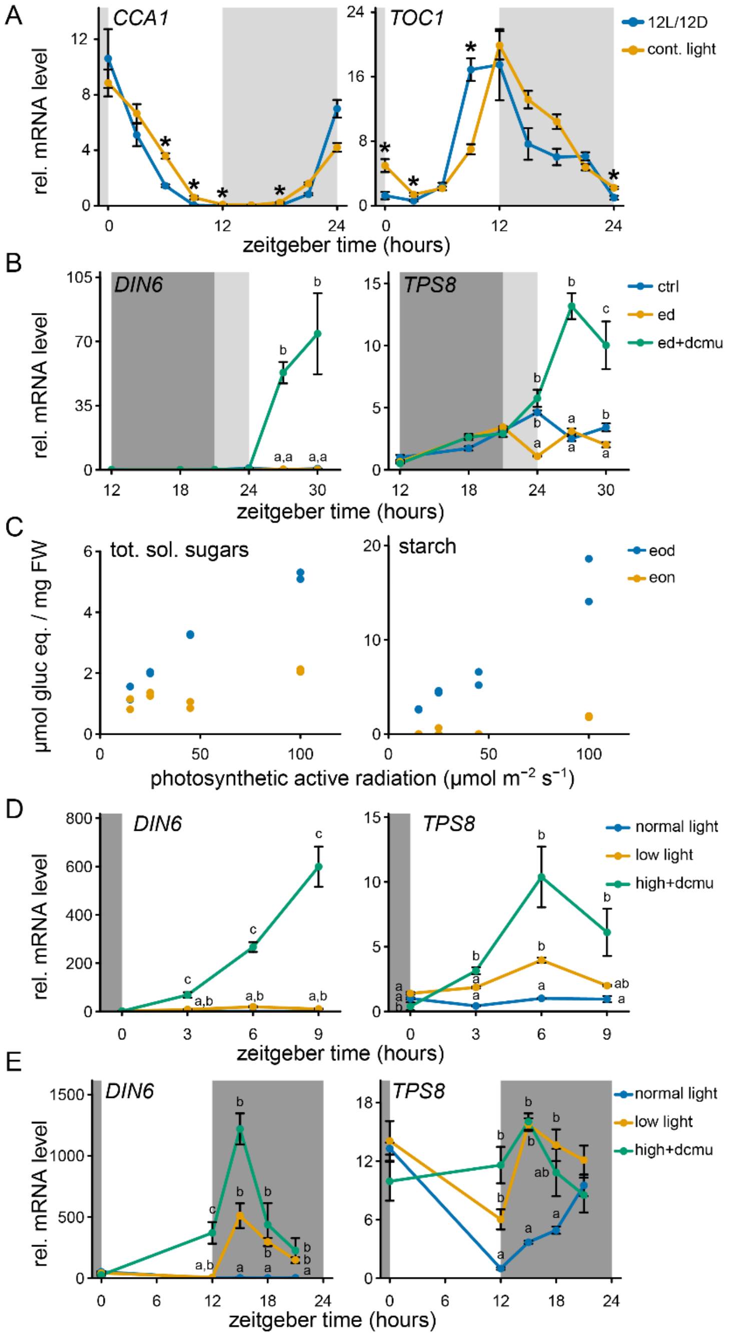


**Supplemental figure 2. The effect of treatments on the clock, sugar dynamics and starvation responses**

**A.** The behaviour of two circadian clock genes (*CIRCADIAN CLOCK ASSOCIATED1 amd TIMING OF CAB1*) during free running cycles as in figure 1A. Light grey boxes represent subjective night. Sampling started second day after starting continuous light conditions. **B.** Effect of a 3 hour early start of the night on carbon starvation marker genes (*DARK INDUCED6* and *TREHALOSE PHOSHATE SYNTHASE S8*). Dark and light grey represent night and the subjective night, respectively. **C.** Carbon status when confronted by a day of reduced light availability. Measurements were taken directly after the low light day (eod, ZT12) and the after the subsequent night (eon, ZT24). **D.** and **E.** The effect of reduced light availability or DCMU on daytime or subsequent nighttime transcript abundance of carb starvation responsive genes. DCMU was applied at the start of the preceding night. Dark grey and light grey boxes represent the night and subjective night respectively. All data are from ~10 leaf stage soil grown Col-0 rosette. Mean ± sem are shown, n=4 (A, B, D and E). Asterisks (planned comparisons) and letters (Tukey HSD) represent statistically significant difference at specific time points $p < 0.05$).

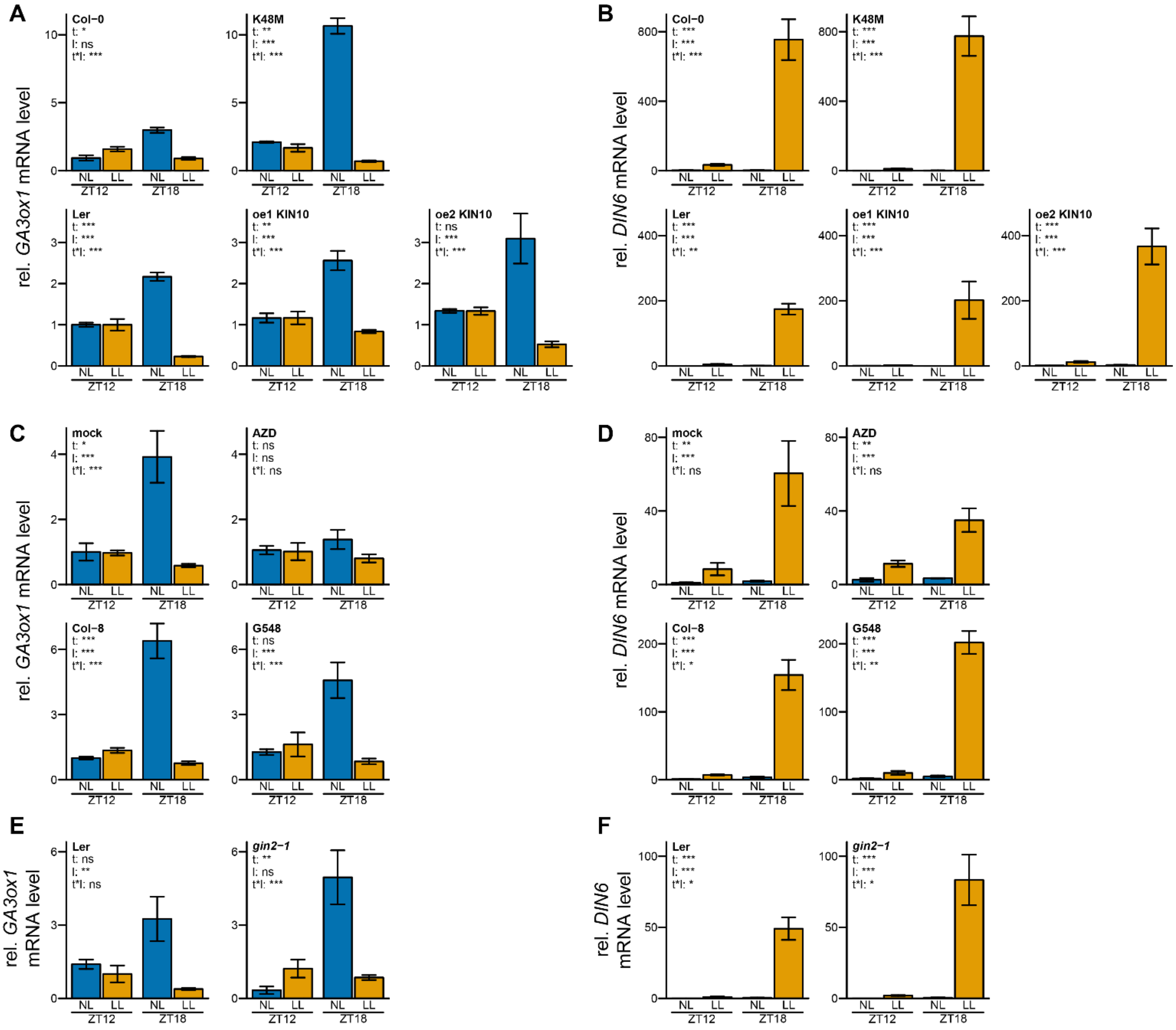


**Supplemental Figure 3. The relevance of sugar/energy signalling pathways on *GA3ox1* expression**

The role of three main energy signalling pathways in the regulation of *GA3ox1* and *DIN6* at the end of the day (ZT12) and during the subsequent night (ZT18), both after control (NL) and low (LL) light levels. **A.** and **B.** K48M: overexpresses the dominant inactive form of SNF1 RELATED PROTEIN KINASE 1 (SnRK1.1/KIN10), where lysine residue 48 is mutated (22). Independent overexpressors 1 and 2 (oe1/2) of KIN10 express the wildtype allele, in Ler background. **C.** and **D.** Manipulation of TARGET OF RAPAMYCIN (TOR) kinase pathway. Mock treatment (DMSO 0.01 %) or AZD8055 (AZD, 30 µM) was applied to Col-0 at 6 am, 2 hours prior the start of the day (ZT0, 8 am). AZD blocks TOR kinase activity. G548 overexpresses TOR in a Col-8 background (23). **E.** and **F.** *Glucose insensitive2-1* (*gin2-1*) and corresponding Ler background (24). Mean ± sem is shown (N=3-5). Asterisks indicate the significance of effects in a 2-way ANOVA. * P < 0.05; ** P < 0.01; *** P < 0.001. t – effect of time (ZT12, ZT18), l – effect of light (NL, LL), t*l, interaction of time and light.

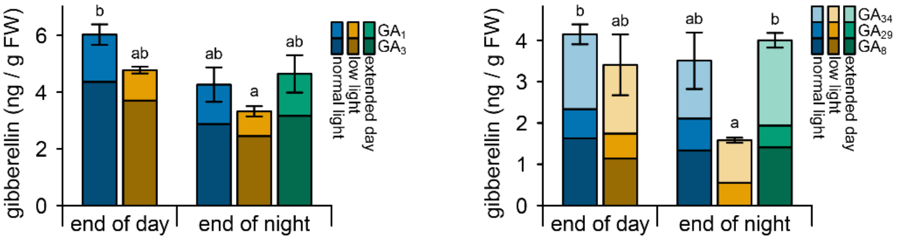


**Supplemental Figure 4. Behaviour of additional GA metabolites over time and during low light conditions**

Abundance of gibberellins, directly following a control (normal light) or low light day (end of day, ZT12) and after the subsequent night (end of night, ZT24). Extended day (under control light conditions) represents a 3 hour delay in the start of the night as in Figure 2C. Data are mean, sem of the sum of compounds is shown. Letters indicate significant differences ($P < 0.05$, Tukey HSD, N=3). $GA_1$ and $GA_3$ are bioactive gibberellins, but with a low affinity for the gibberellin receptor GID1. $GA_{34}$, $GA_{29}$ and $GA_8$ are gibberellins that were made inactive by GA2ox.

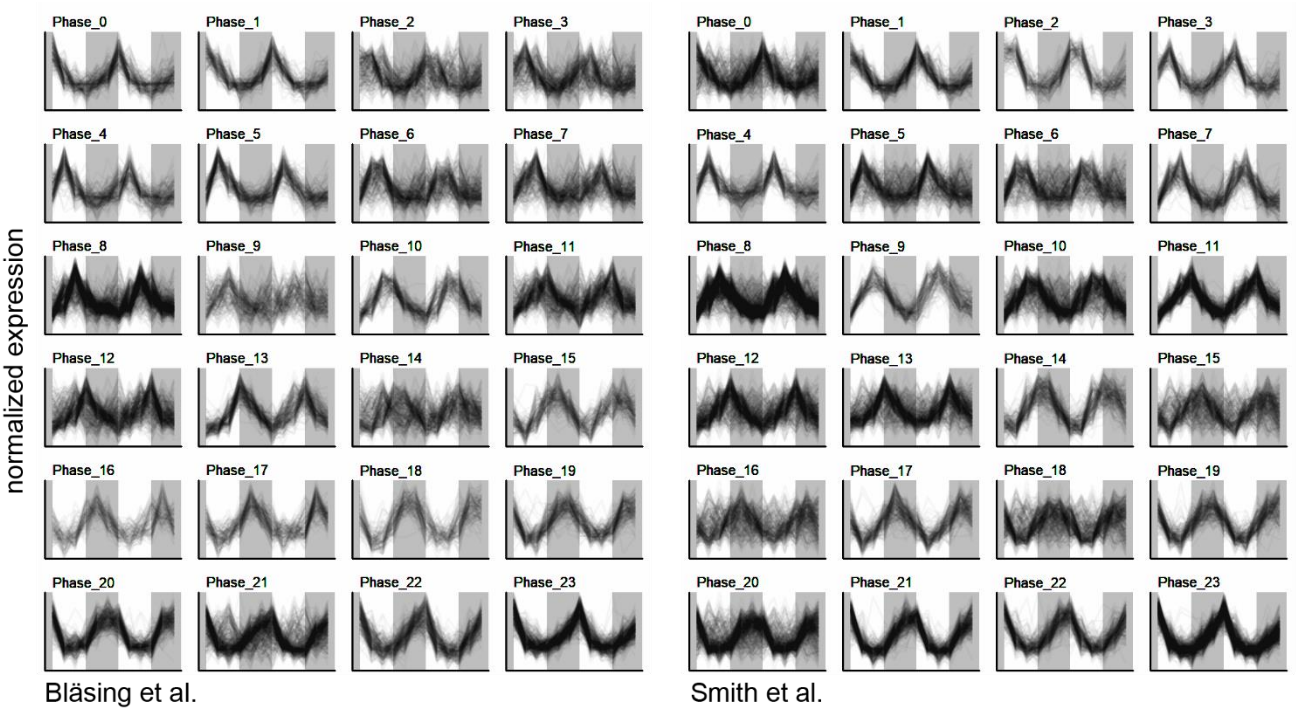


**Supplemental Figure 5. Rhythmic expression of gene sets tested for gibberellin enrichment.**

Rhythmic genes in two independent datasets (20-21, 47). Gene sets for each phase were pooled with a sliding window approach. Window size: 3 hours / 3 timepoints. Stepsize: 1 hour / 1 time point. Unpooled data are shown.

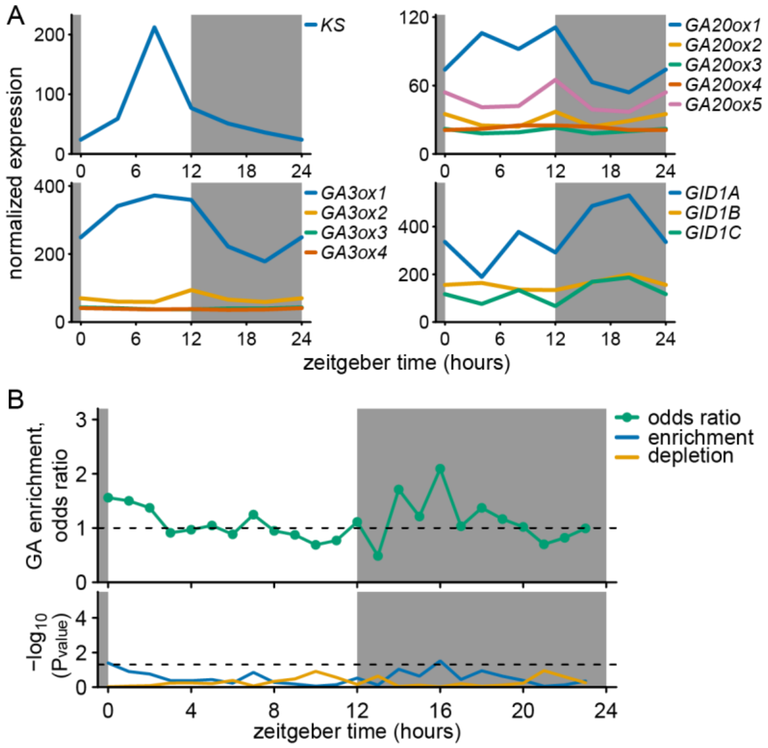


**Supplemental Figure 6. GA biosynthesis gene behaviour and GA responsive genes in *pgm***

**A.** The mRNA abundance levels of key rhythmic gibberellin biosynthetic genes and of the gibberellin receptor *GID1*. The public data is from soil grown starch mutant *pgm* at a light intensity of 130 µmol/m$^2$ (21). For clarity, the unsampled ZT24 is represented by values from ZT0. **B.** Relative enrichment (odds ratio > 1) or depletion (odds ratio < 1) of GA responsive genes among gene sets that have rhythmic behaviour peaking at specific times of the day in the *pgm* mutant (21,47). GA responsive genes were based on GA treated rosettes (25). Below is the statistical significance of depletion or enrichment of GA responsive genes (Hypergeometric distribution). Dashed line indicates P = 0.05.

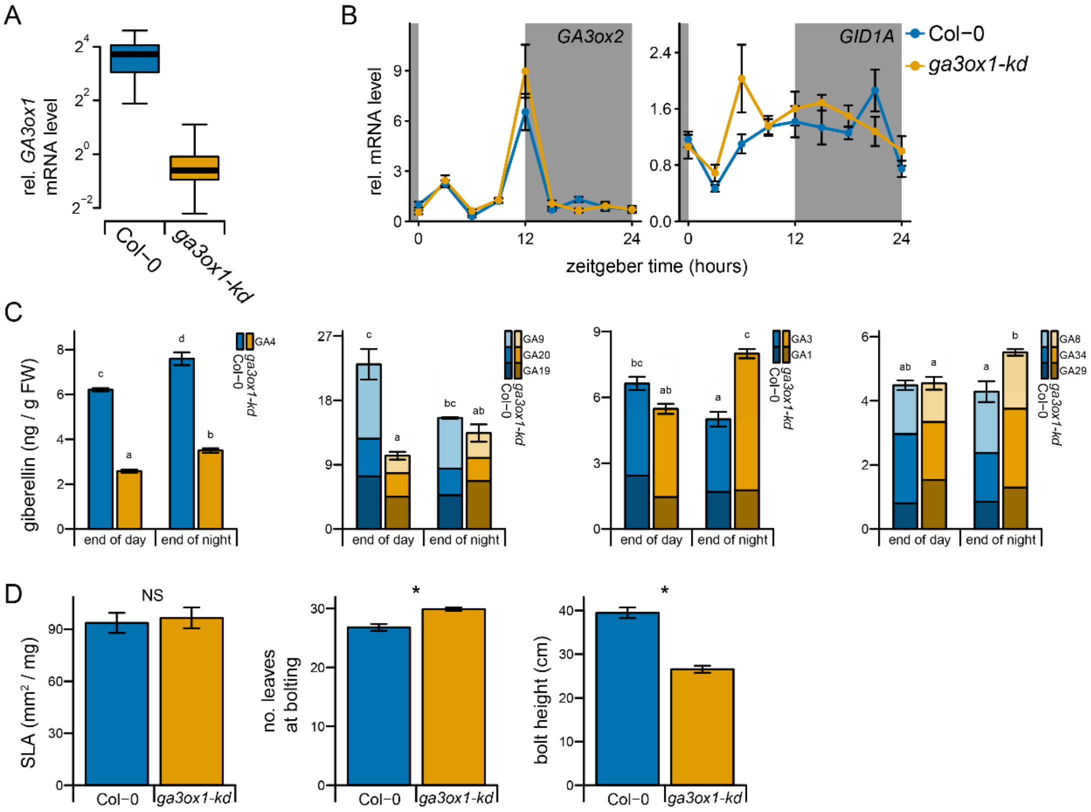


**Supplemental Figure 7. General characterization of *ga3ox1***

**A.** Difference in transcript abundance on a log scale over the entire day night cycle between wild type and *ga3ox1-kd*, mean ± sem (N=4). **B.** mRNA abundance of key genes involved in GA metabolism or signalling during the day night cycle. **C.** Gibberellin levels in Col-0 and the mutant under control conditions. Mean ± sem (N=3) are shown. From left to right, main bioactive $GA_4$, precursors, low affinity bioactive GAs and inactivated GA. **D.** Morphological and developmental traits of *ga3ox1-kd* compared to Col-0. Mean ± sem are shown (N=10). SLA is specific leaf area. SLA is based on total leafblade area and corresponding weight of plants at the 11 leaf stage. Asterisks and letters (t-test / Tukey HSD ) represent statistically significant difference at specific time points $p < 0.05$).

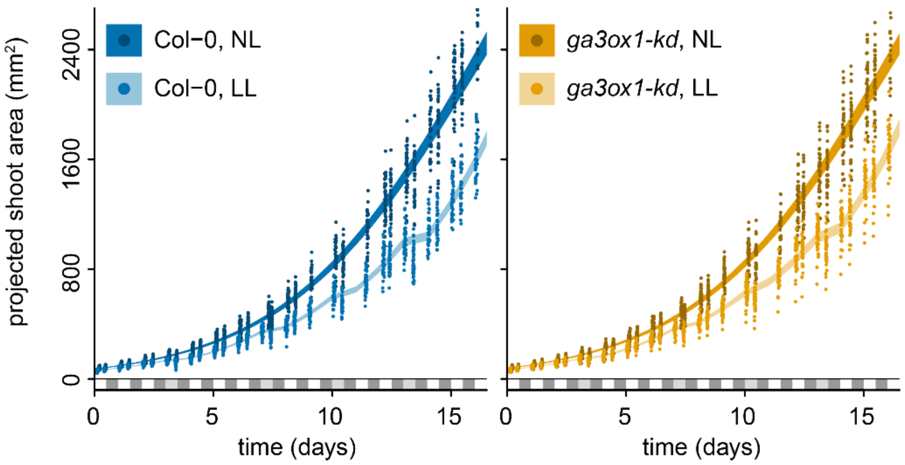


**Supplemental Figure 8. carbon and GA dependent growth under constant and variable conditions**

**A.** Points indicate individual measurement. The shaded area represents the 95 % confidence interval of the mean, as estimated by the growth curves to individual plants. N=40. The white and dark grey boxes represent the day night cycle. Light grey at day 4, 8, 11 and 14 indicate days with the low light treatment.

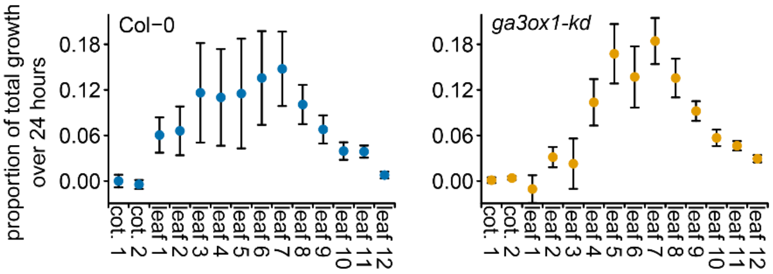


**Supplemental Figure 9. Contribution of different leaves to total rosette growth**

Area of individual leaves were determined by dissecting rosettes either at ZT0 or ZT24. Growth rates are based on the increase in leaf area. For most plants, leaf 12 was newly formed in the 24 hour period. Values are mean ± sem (n=22 for Col-0; n=32 for *ga3ox1-kd*). Cot. is cotyledon.

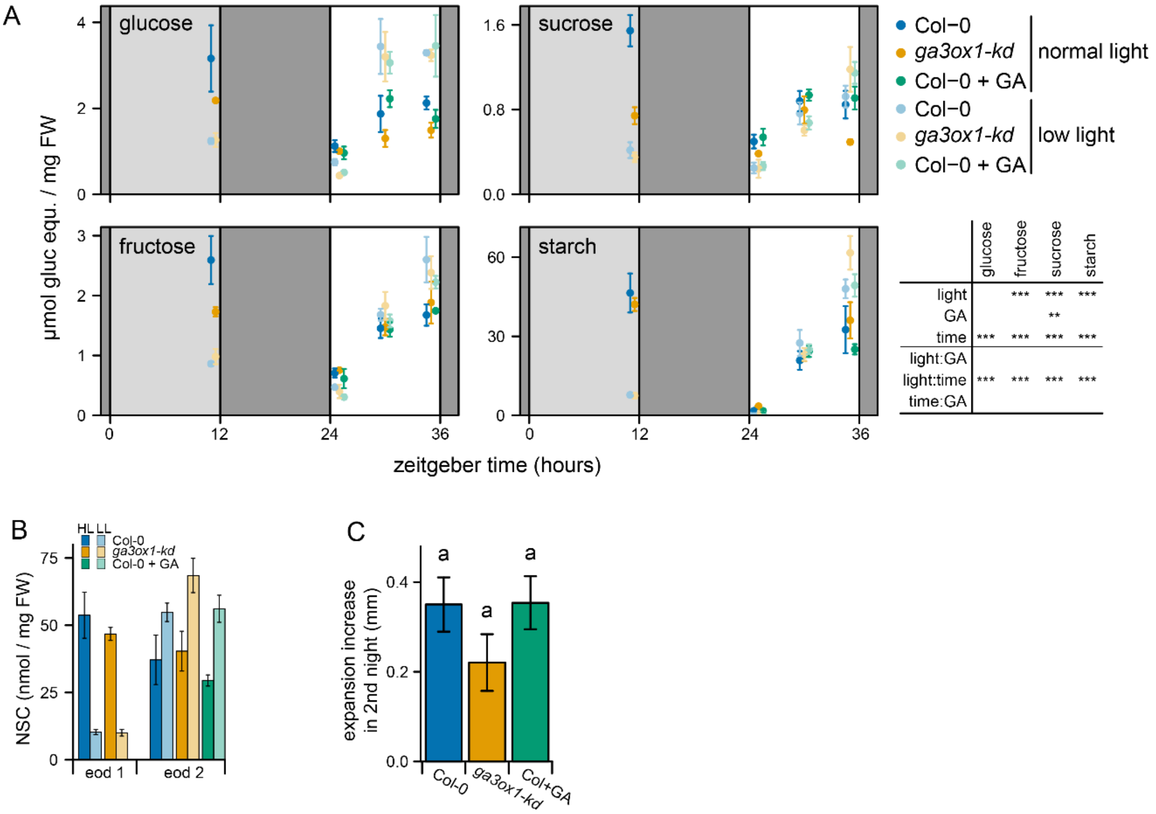


**Supplemental Figure 10. Non-structural carbohydrates during and after a low-light day**

Experiment performed as in Figure 6F. **A.** Dark and light grey boxes indicate the night and the low light treatment respectively. Gibberellin/mock was applied at ZT12. Data are mean ± sem, N=3. Bottom right results of three-way ANOVA without tree way interaction term, for each compound. Light effect (normal light, low light), GA effect (Col-0, *ga3ox1-ks*, GA addition), time effect (ZT12, ZT24, ZT30, ZT36). * P < 0.05, ** P < 0.01, *** P < 0.001. **B.** Total non-structural carbohydrates in glucose equivalents (NSC: starch, sucrose, glucose and fructose) at the end of a low light day (eod 1, ZT12) and at the end of the subsequent day (eod 2, ZT36), same data as in A. mean ± sem, N=3. **C.** Expansion increase compare to normal light (NL) plants during the second night after a low light day (ZT36 to ZT48), mean ± sem (N=25). Col+GA is Col-0 + GA under HL compared to Col-0 high light + mock. Letters indicate significant differences (P < 0.05, Tukey HSD). NL: control normal light conditions, LL: low light conditions.

**Supplemental table 1.** Primers used for RT-qPCR

| Gene | AGI code | Direction | Sequence 5' -> 3' |
|---|---|---|---|
| *TIP4-like* (Reference gene) | AT4G34270 | Forward | AGTGAGAGTCATGCCAAGCT |
| | | Reverse | CAGTTGGTGCCTCATCTTCG |
| Unknown protein (Reference gene) | AT1G13320 | Forward | GTGAGGGAGAAAGCTGTGGA |
| | | Reverse | AGCCAGAGGAGTGAAATGCT |
| *KAURENE SYNTHASE* | AT1G79460 | Forward | GTGGAGCTTTCTGTTTCGGC |
| | | Reverse | ACTGTGGGAAAAGTGGAGCA |
| *GA20OX1* | AT4G25420 | Forward | GGTTTCTTCCTCGTGGTCAA |
| | | Reverse | TTTCGGAGAGAGGCATATCAA |
| *GA3OX1* | AT1G15550 | Forward | CGATTTCCGTAAACTTTGGC |
| | | Reverse | GACTGGCCCATTCAATGTCT |
| *GA3OX2* | AT1G80340 | Forward | CCCAGCCACCACCTCAAATA |
| | | Reverse | ACTCCCAGTGAACCTAATGCG |
| *GID1A* | AT3G05120 | Forward | GCATCCAGCGTGTAATCCGT |
| | | Reverse | CGCGACAACCACAAGACTCT |
| *TOC1* | AT5G61380 | Forward | TGATCTTCCCAATGGCTAAGG |
| | | Reverse | ACTTCGTCTTGCCTCGACAT |
| *CCA1* | AT2G46830 | Forward | ATCCTCGAAAGACGGGAAGT |
| | | Reverse | TCAGGCTTTGATTGTTGTCG |
| *DIN6* | AT3G47340 | Forward | AAGGTGCGGACGAGATCTTTGG |
| | | Reverse | ACTTGTGAAGAGCCTTGATCTTGC |
| *TPS8* | AT1G70290 | Forward | GTGGTTGTCAAGAGAGGTCAACAC |
| | | Reverse | AGCTAGACCTTTGCTTACACCTTG |
